# Global trends in antimicrobial resistance on organic and conventional farms

**DOI:** 10.1101/2023.04.07.536071

**Authors:** Eldon O. Ager, Tamilie Carvalho, Erin Silva, Steven C. Ricke, Jessica L. Hite

## Abstract

Various stewardship policies, regulations, and voluntary bans have focused on protecting antimicrobials by limiting their use in livestock. These efforts ignited management shifts ranging from largely nominal (e.g., drugs banned for use as ‘growth promoters’ were reclassified as ‘prophylactic’ drugs) to organic farming, which drastically reduces or eliminates use of antimicrobials. Understanding how these farming practices influence the prevalence of antimicrobial resistance in livestock carries important implications for policy makers, public health officials, and farm managers. Here, we reviewed studies spanning the last 20 years to ask if the most stringent effort to reduce antimicrobial use in livestock — organic farming — results in notable reductions in the prevalence of antimicrobial resistance across broad scale geographic ranges, pathogens, and livestock hosts. Our results validate organic farming in reducing the prevalence of antimicrobial resistance (AMR) by ∼31.2%,∼26.9%, ∼28.2%,∼42.9 and ∼36.2% in cattle, chicken, environment, pigs and turkey respectively while also revealing significant variation in the strength of this reduction across contexts. Given that our results join others indicating that AMR is increasing across all types of farms, our results highlight areas where organic farming has been most effective and may provide economical and scalable solutions for farmers.

## INTRODUCTION

Antimicrobial resistance is an increasingly urgent global health crisis that disproportionately affects developing nations and poor livestock farmers, of which approximately 67% are women. Globally, the majority (∼73%) of all antibiotics are used in animals raised for food ^1^ with over 45 mg/population correction unit (PCU) in cattle, 148 mg/PCU and 172 mg/PCU in chickens and pigs, respectively ^2^. The use (and overuse) of antimicrobials in livestock is generally linked with the rise of drug-resistant infections — both in animals and humans. Yet, the specific suite of farming practices driving this broad pattern remains poorly resolved. Thus, there is an urgent need to identify specific farming practices that promote or hinder drug resistance and to understand whether the rates and patterns of drug resistance are contingent on specific geographic regions or host types.

As a first step in this endeavor, we focus on synthesizing global patterns of antimicrobial resistance across livestock hosts on both organic and conventional farms. Why focus on organic farming? Organic farming leads the agricultural industry in efforts to reduce the overuse of antimicrobials in livestock by integrating cultural, biological, nutritional, and mechanical methods to ensure environmentally safe and residue-free foods, along with improved animal welfare standards ^3–6^. Organic production also provides animals with a more spacious and enriched environment, access to an outdoor range, limited group sizes, and other preconditions, all of which reduce the need for medications, including antimicrobials .

Globally, ‘antibiotic-free’ livestock products are in high demand ^7^ leading the shift towards organic agriculture. In the U.S., organic is the fastest-growing sector of the food industry, with organic meat and poultry sales increasing by 2.5% and 4.7% respectively in 2021 ^8^. These trends in organic farming carry important implications for disease outbreaks and the spread of antimicrobial resistance. Indeed, mounting evidence suggests that organic farming practices can reduce the occurrence of pathogenic outbreaks and, therefore, the presence of genes that carry and spread antimicrobial resistance ^9–11^. These patterns suggest a potential win-win for animal welfare, conservation, and public health that move far beyond stewardship policies that vary widely across geographic regions and can unintentionally drive an increase in antimicrobial use.

For example, the European Union (EU) banned the use of antimicrobial growth promoters (AGP) in livestock production systems due to AMR in 2006 ^12^ while the United States (US) implemented a sweeping ban on the use of AGPs in 2017 ^13^. Unfortunately, countries across Europe reported an *increase* in the use of antimicrobials for prophylactic purposes after the ban of AGPs. These well-intentioned policy changes backfired due, in part, to a loophole in naming conventions, as well as a lack of practical and scalable alternative solutions for farmers who depend on antimicrobials to prevent outbreaks and maintain herd health. This pattern underscores the complex challenges of reducing the use of antimicrobials in livestock.

Organic farming practices offer a promising alternative to imposing top-down restrictions on drug use in livestock. While the goals of organic farming practices are multifaceted, helping to combat the antimicrobial resistance crisis remains a central goal for this burgeoning industry. A better understanding of how organic farming practices maintain animal health without reliance on antimicrobials could identify novel approaches for conventional farms. However, our understanding of the extent to which organic farming practices can successfully reduce the emergence and spread of AMR remains fragmentary — and hotly debated ^14–17^. An important research objective, therefore, is to understand how broad-scale patterns of antimicrobial resistance vary across organic and conventional farms.

We conducted a large-scale literature review to test the long-standing hypothesis that organic farming reduces the prevalence of antimicrobial resistance. We found support for this hypothesis, as well as clear evidence that the decline in drug resistance varies across contexts.

Taking these findings together, our study underscores the potential for organic farming practices to strongly limit antimicrobial resistance while highlighting the possibility that environmental contamination may undermine these efforts. These findings further emphasize an increasing temporal trend in antimicrobial resistance across both organic and conventional farms, even for drugs considered critically important for human medicine, suggesting that current policies to protect these powerful biomedical interventions fall short.

## METHODS

### Literature search strategy

We conducted literature searches for studies published between 2000-2022 using three electronic databases (PubMed, Web of Science, and PubAg). We used the following search terms, which we modified slightly for each database. Note, for brevity, we show abbreviated terms (e.g., “livestock names” reflects individual searches for sheep, goats, chickens, etc., and the name of the pathogen reflects individual searches for each pathogen ‒ i.e. *Campylobacter*, *E. coli*, *Salmonella* etc. In the PubMed search, for example, used the following terms: livestock name AND product OR livestock production OR livestock farm OR name of pathogen OR antimicrobial AND resistan OR agriculture OR conventional OR organic AND agriculture. The full search terms are provided in the Supplementary Materials.

Finally, we screened references from other literature reviews ^18–20^ and all other studies were screened for inclusion. Our search produced 1,836 hits (Fig.1A). We used the term “organic farm” to refer to farms that either eliminate or drastically restrict the use of antimicrobials for the treatment of animals, while “conventional farm” refers to farms that use antimicrobials for the treatment of animals ^21^.

**Figure 1.**
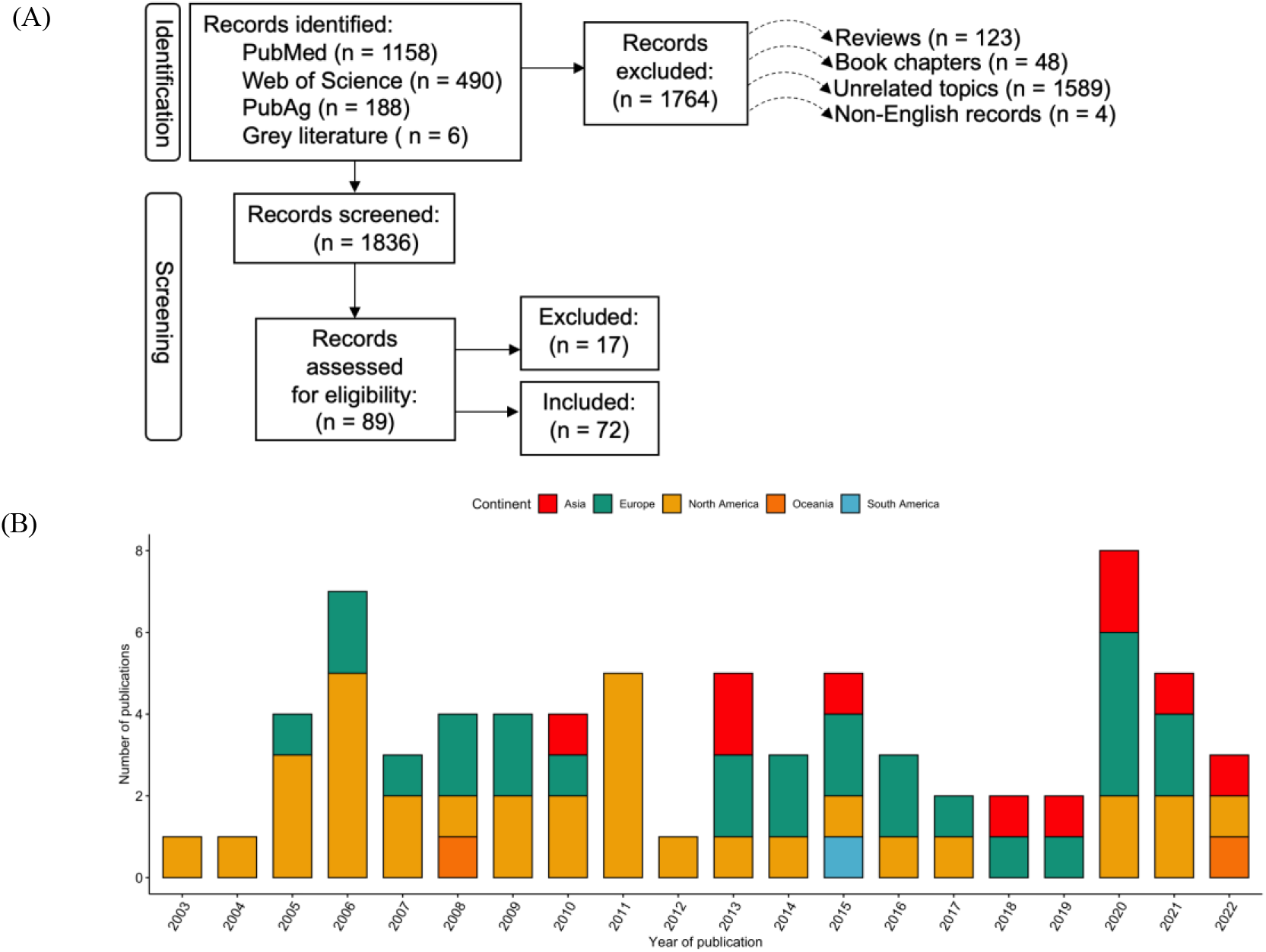
(A) PRISMA flow diagram showing the number of studies included in the review. (B) Overview of studies and geographies covered in this review. We focused on 72 studies published between 2000 and 2022 that included point prevalence surveys of antimicrobial resistance on organic and conventional farms. Of these, 46% were in North America (n = 33), 36% in Europe (n=26), and 14% in Asia (n = 10), while Oceania and South America contributed to 3% (n = 2) and 1% (n=1), respectively.

### Study selection

We reviewed all full English-written articles that directly compared patterns of antimicrobial resistance (AMR) in poultry, chicken, turkey, cattle, and pigs, and in environmental samples from organic and conventional farms in a given geographic region. These included paired studies (conventional vs. organic farms) and unpaired studies focusing on either organic or conventional farms. Studies were only included if they clearly described the farms as organic or conventional, presented antimicrobial resistance data, and clearly defined the study location. After screening the reference lists, we excluded reviews, non-English records, unrelated topics, and book chapters (Fig. 1A). Following the search, all records were exported to Endnote’s web citation manager ^22^, where duplicates were removed. The records were then exported to a spreadsheet and organized by title, doi, authors, journal, year of publication, and abstract. Finally, the titles and abstracts were screened against the inclusion criteria.

The studies that met the eligibility criteria were retrieved in full text and were thoroughly reviewed. Seventy-two studies met our inclusion criteria (Fig. 1A). We attributed the reduction in sample size to the fact that our search terms covered general antimicrobial resistance and antimicrobial susceptibility topics. As a way of assessing data quality in our review, we excluded records that did not clearly identify farm types as organic or conventional, gave no geographic information on study location, provided unclear resistance rates, or involved imported products. Data extraction results were stratified according to country name, antimicrobial resistance results, farm type, and pathogens.

### Statistical analysis

All data analyses were conducted in R version 4.2.0 ^23^ and QGIS version 3.24.0-Tisler ^24^. We tested whether antimicrobial resistance differed significantly between organic and conventional farms. The level of significance was set at p<0.05. We used generalized linear mixed-effects models (GLMM) with binomial distributions and log-link function. The response variable was percentage resistance; farm type and host type were set as fixed effects; farm ID, country, continent, antimicrobial compound, antimicrobial, class, pathogen, and year of sampling were set as random effects. We used restricted maximum likelihood using R’s nlme (package RELM ^23, 25^. Following Burnham and Anderson (2004) ^26^, we compared candidate models using Akaike’s information criterion (AIC) and ΔAIC to identify the best model (the model with substantial support). We then evaluated the factors in those models with visual diagnostics, including Turkey-Anscombe plots, quantile-quantile plots, and residual-versus-predictor plots ^27^ (see Supplementary Materials). To report resistance in foodborne pathogens, we calculated the pooled prevalence of resistance from each pathogen-drug resistance rate to report AMR in foodborne pathogens as described by ^1, 28^.We used the percentage change formulae below to calculate the percentage antimicrobial resistance change between organic and conventional farms across hosts.

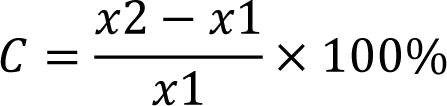

𝐶 = percentage resistance change

𝑥1 = organic farm resistance prevalence

𝑥2 = conventional farm resistance prevalence

## RESULTS

We identified 1,833 unique academic studies published between 2000 and 2022, as well as six grey literature publications, that reported the point prevalence of antimicrobial resistance. After the references were subjected to relevance screening, 1,744 academic studies and all six grey literature publications were removed. After assessing the remaining 89 references for eligibility, 17 studies were excluded. The systematic review included 72 studies that met the inclusion criteria (Fig. 1A). North America accounted for 46% (n = 33) of the surveys; Europe accounted for 36% (n = 26); 14% of studies (n = 10) were from Asia, while Oceania and South America contributed 3% (n = 2) and 1% (n = 1), respectively (Fig. 1B). All the surveys covered antimicrobials classified as critically important and highly important for human medicine by the World Health Organization.

### Resistance trends across organic and conventional farms

Overall, conventional farms had higher mean antimicrobial resistance (28%) relative to organic farms (18%; *t* = -3.616, *p* = 0.0003). However, between 2001 and 2020, the percentage of antimicrobial resistance increased on both organic and conventional farms. The percentage of AMR on organic farms increased from 10% (CI: 2-18%) to 32% (CI: 23-41%), while the percentage of resistance on conventional farms increased from 18% (CI: 12-28%) to 38% (CI: 33-51%; *t* = *-*2.429, *p* = 0.0215; Fig. 2A and 2B).

**Figure 2.**
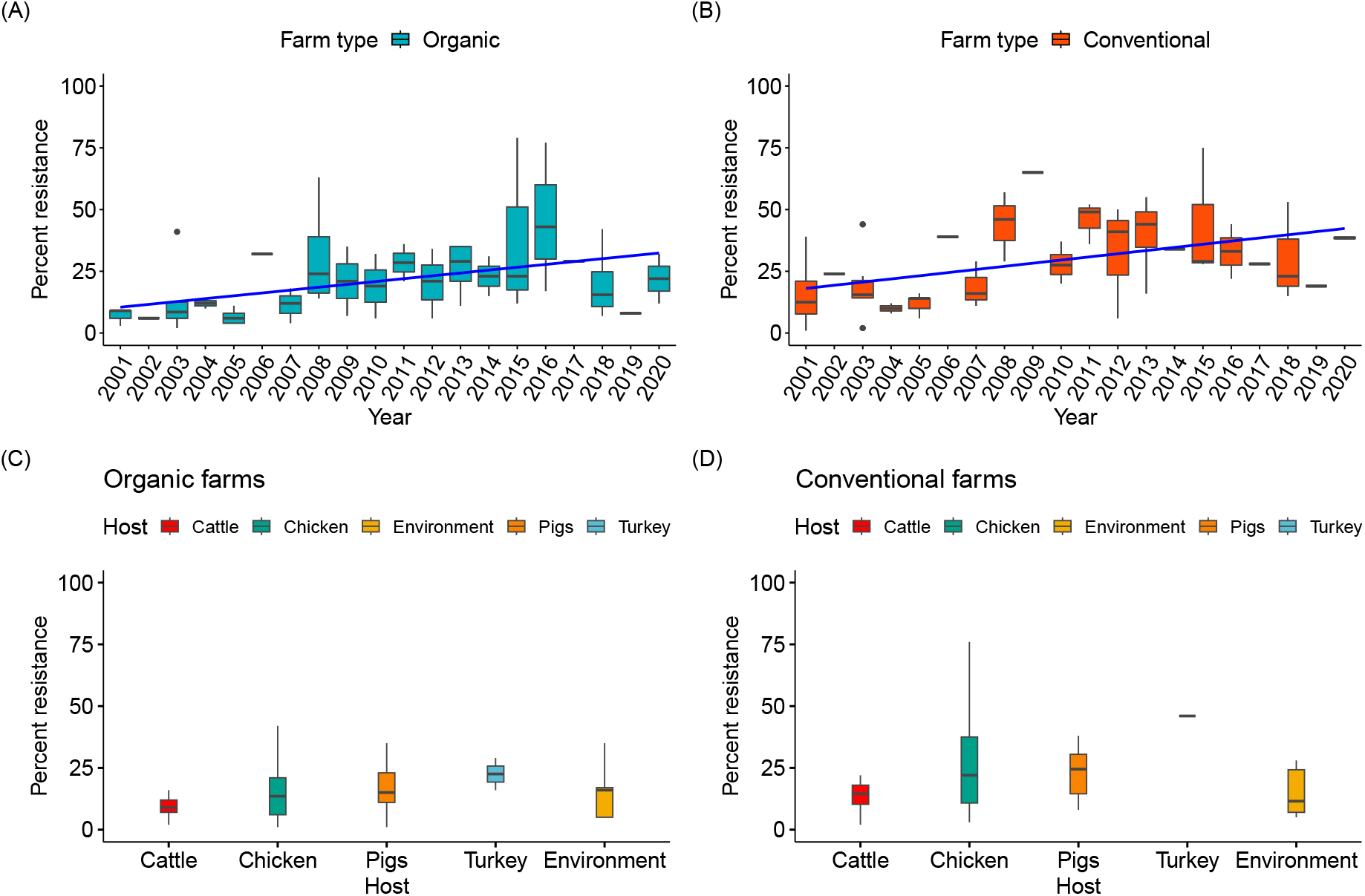
Temporal trends in antimicrobial resistance on organic and conventional farms between 2001 and 2020 and across different sample types. [A-B: (A) Organic farm surveys, n = 56; (B) conventional farm surveys, n = 53]. Twelve surveys were excluded from this analysis because they did not report the study years. Overall, antimicrobial resistance was higher on conventional farms at 28% compared to organic farms at 18%. However, antimicrobial resistance appears to be increasing on both organic and conventional farms. The percentage of antimicrobial resistance in isolates from organic farms increased from 10% (95% CI: 2-18%) to 32% (95 % CI: 23-41%) while on conventional farms the percentage resistance increased from 18% (95% CI: 12-28%) to 42% (95% CI: 33-51%). [C-D: (C) Organic farm isolates, n = 29,417; (D) conventional farm isolates, n = 31,882. Across cattle, chicken, pigs, and turkey isolates, the prevalence of antimicrobial resistance in cattle, chicken, pigs, and turkeys on conventional farms was higher as compared to organic farms. However, antimicrobial resistance was higher in environmental samples collected from organic farms, ostensibly because of prior contamination.

### Host-specific patterns

Resistance patterns were highly variable across different host classes, with overall resistance higher in hosts from conventional farms. Surprisingly, the AMR prevalence in environmental isolates was higher on organic farms (16%) than on conventional farms (11.5%; *t* = -3.576, *p* = 0.0004; Fig. 2C and 2D). However, conventional farms reported a higher AMR prevalence in isolates from cattle, chicken, pigs, and turkey. For instance, in isolates from cattle, the prevalence of AMR was 14.5% on conventional farms and 9% on organic farms (*t* = 6.826, *p* = 0.000; Fig 2C and 2D). For chicken isolates, AMR prevalence was higher on conventional farms (22%) compared to organic farms (13.5%; *t* = -2.401, *p* = 0.0165). Similar trends were reported for other hosts. For example, resistance was higher on conventional pig farms (24.5 %) than on organic farms (15 %; *t* = -6.782, *p* = 0.00; Fig. 2C and 2D). Conventional turkey farms reported a significantly higher AMR prevalence (46%) as compared to organic farms (22.5%; *t* = -2.808, *p* = 0.005; Fig. 2C and 2D).

### Geographical distribution of AMR across regions

The prevalence of antimicrobial resistance varied across geographic regions by host and farm type. Some regions reported high AMR prevalence in environmental isolates on conventional farms as compared to organic farms. For instance, some parts of the USA reported AMR prevalence of 64% (n = 135) and 35% (n = 135) in environmental isolates on conventional and organic farms, respectively. As expected, countries with low antimicrobial usage in food animals, like Sweden and New Zealand, reported low AMR prevalence in the environment. For instance, Sweden reported 5% (n = 725) AMR prevalence on both organic and conventional farms, while New Zealand reported 5% AMR prevalence on conventional farms (n = 814) and 3.8% on organic farms (n = 814; Fig. 3).

**Figure 3.**
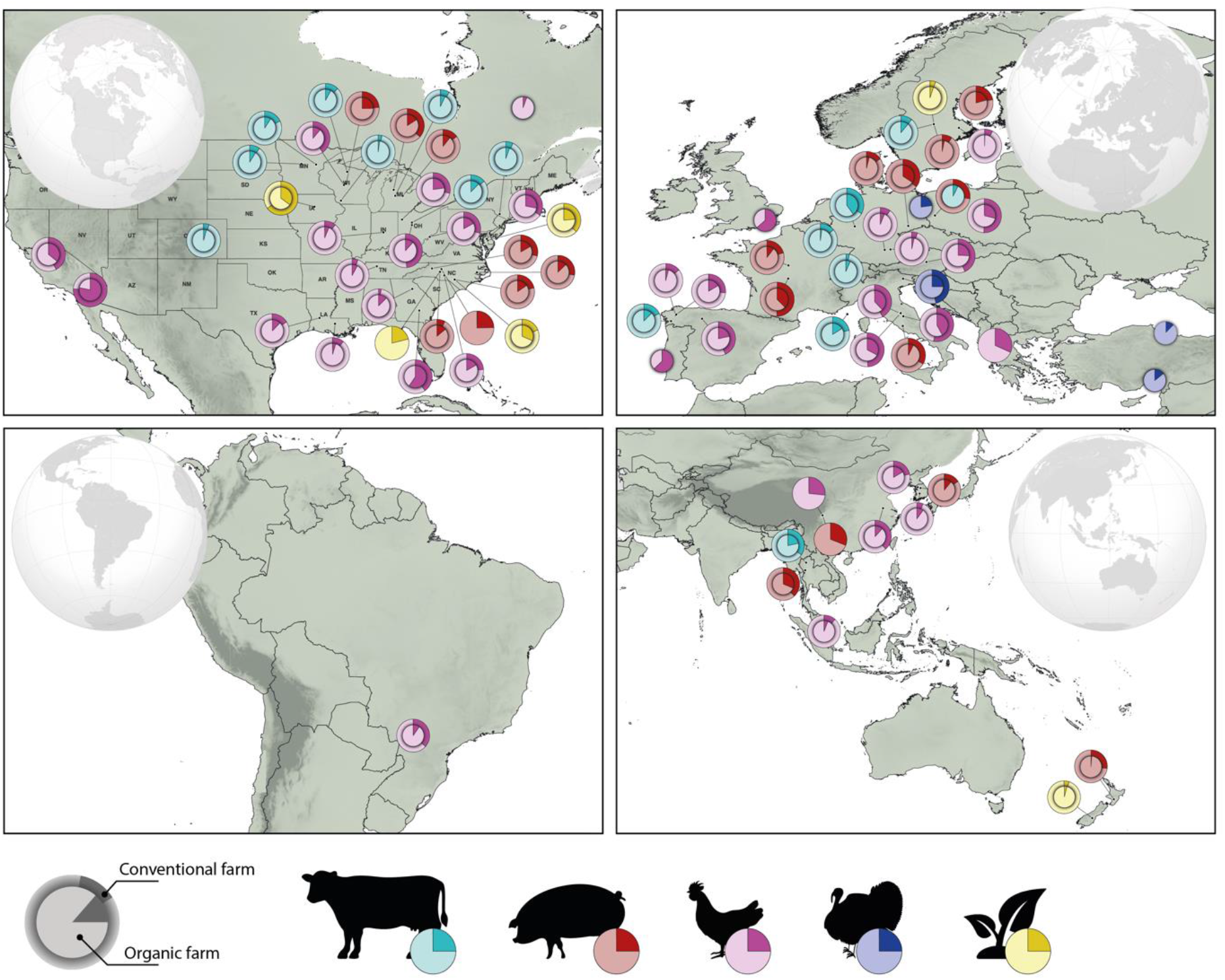
Global patterns of antimicrobial resistance in isolates collected from organic and conventional farms. Studies spanned four hosts (bovine, porcine, chicken, and turkey) and environmental samples collected from conventional and organic farms throughout North America, Europe, Asia, Oceania, and South America. Pie charts show the prevalence of antimicrobial resistance on conventional farms (outer pie chart, *n* = 66) relative to their organic counterparts (inner pie chart, *n* = 69). Geographic regions with a single pie (i.e., outer pie only), represent areas lacking data from organic farms.

Surprisingly, AMR in isolates from chicken was higher on organic farms than on conventional farms in two different regions of Georgia, USA: 40% on conventional farms (n = 60) and 59% on organic farms (n = 60) in one region and 3% on conventional farms (n = 40) and 13% on organic (n = 40) farms in another region. In California, USA, prevalence of AMR in chicken isolates was significantly higher on both farm types: 78% on organic farms (n = 132) and 75% on conventional farms (n = 132). Organic farms in the United Kingdom and Portugal also reported higher AMR prevalence in chickens, 63% (n = 30) and 61% (n = 33), respectively (Fig. 3).

### Patterns in foodborne pathogens

Our study examined patterns of AMR in *Escherichia coli*, *Salmonella spp*, *Campylobacter spp*, and *Staphylococcus aureus*, covering 29,417 isolates from organic farms and 31,882 isolates from conventional farms in Asia (n = 1,164), Oceania (n = 3,085), Europe (n = 11,759), South America (n = 312), and North America (n = 44,979). We found a higher prevalence of AMR in *E.coli* isolates against commonly used antibiotics such as penicillin, tetracycline, and amoxicillin-clavulanic acid across different regions. For instance, Asia reported high AMR prevalence against amoxicillin-clavulanic acid at 98% (CI:95-100%) on both organic (n = 105 isolates) and conventional farms (n = 110 isolates; Fig. 4A). Moreover, a higher AMR prevalence in *E. coli* and *salmonella* was reported against other critically important antimicrobials like erythromycin on both conventional and organic farms in North America and Asia. The prevalence of resistance to erythromycin in *E.coli* was 77% (CI: 75-80%) and 60% (CI: 58-63%) on conventional and organic farms in North America, respectively (n = 785; Fig. 4C), while the resistance prevalence of ampicillin and nalidixic acid were high on conventional farms as compared to organic farms in South America (Fig.4D). As expected, we reported low AMR prevalence to *E.coli* in Oceania as compared to other regions on both organic and conventional farms (Fig.4E). We also reported a higher resistance prevalence against erythromycin in *salmonella* isolates was 100% (n = 171) on conventional farms and 98% (CI: 96-100%) (n = 165) on organic farms in Asia (Fig. 5A) while AMR prevalence against tetracycline in *salmonella* was 24%, CI:18-26% in North America as compared to Asia 18% CI:15-22% (Fig. 5A and 5B).

**Figure 4.**
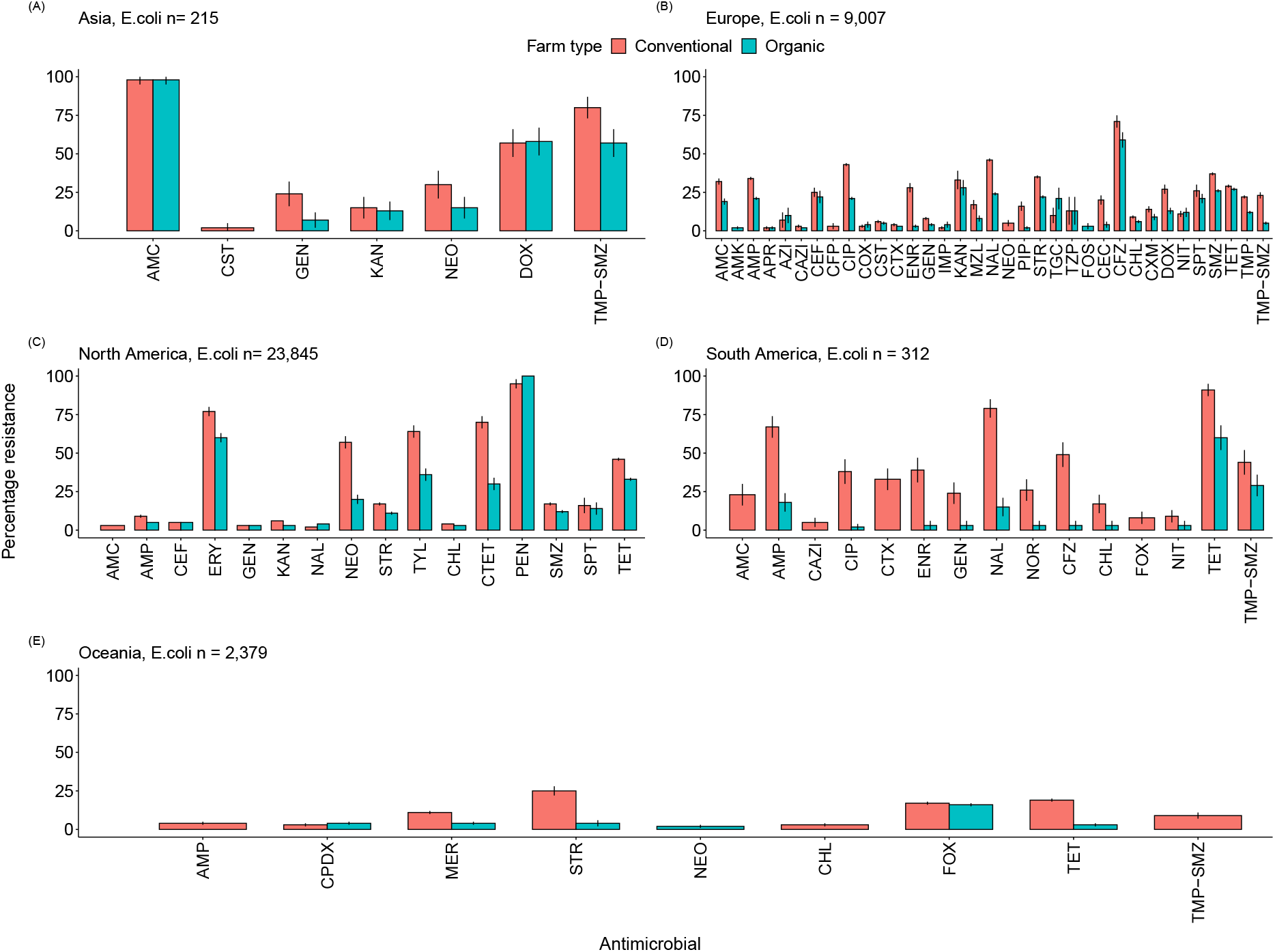
Patterns of antimicrobial resistance in *E. coli.* The percentage of antimicrobial resistance is shown for the number of isolates (n) examined on organic and conventional farms in each geographic region. A) Asia, n = 215, B) Europe, n = 9007, C) North America, n = 23845, D) South America, n = 312 and E) Oceania, n = 2379. The error bars represent the 95% proportion confidence interval. For drug acronyms, see Table1. The grey shading indicates antimicrobials classified as critically important (while the unshaded region indicates highly important antimicrobials).

**Figure 5.**
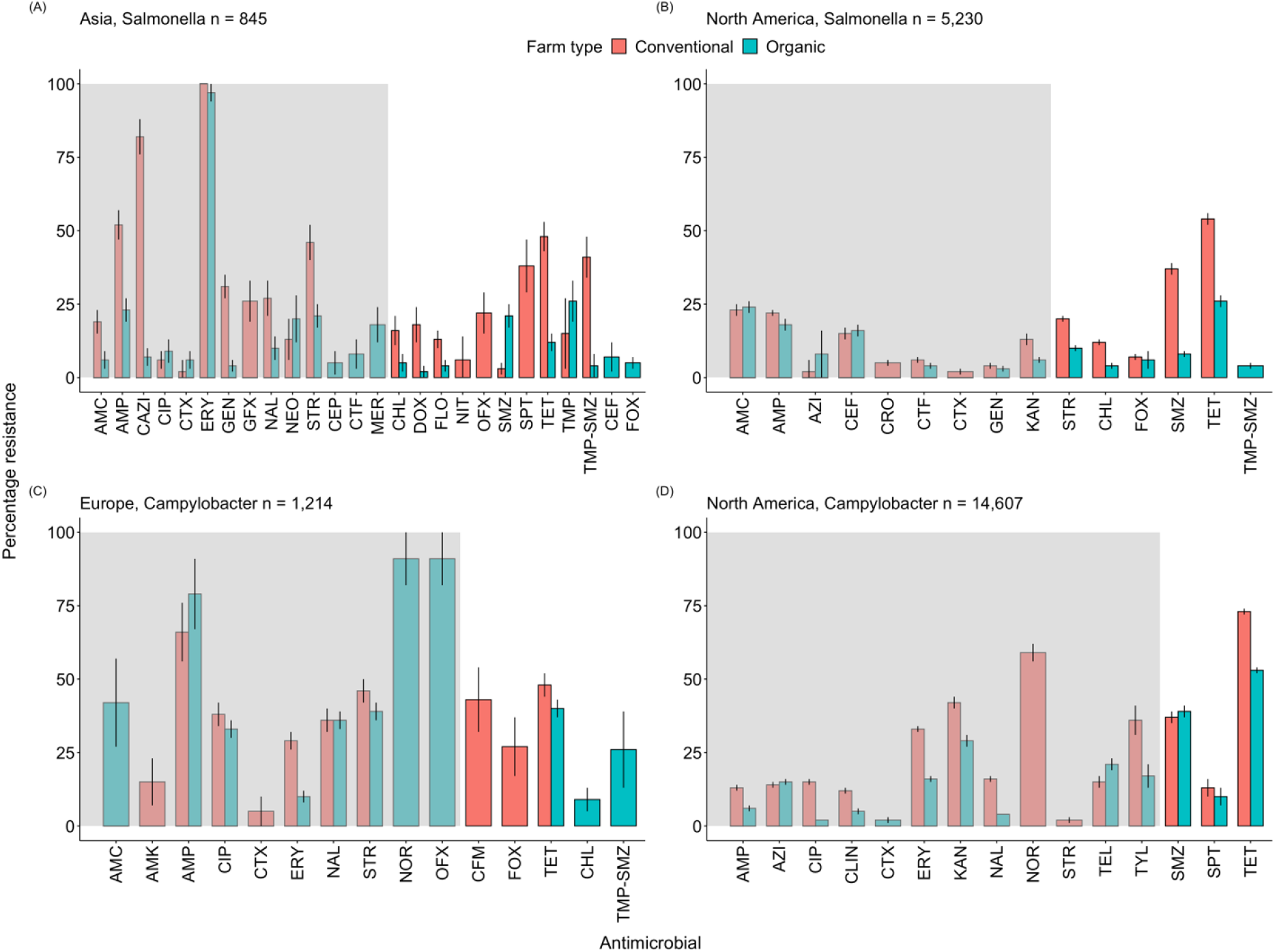
Patterns of antimicrobial resistance in *Salmonella* and *Campylobacter.* The percentage of antimicrobial resistance is shown for the number of isolates (n) examined on organic and conventional farms in each geographic region. A) Asia, *salmonella*, n = 845, B) North America, *salmonella,* n = 5230, C) Europe, *campylobacter,* n = 1214 and D) North America, *campylobacter,* n = 14607. The error bars represent the 95% proportion confidence interval. For drug acronyms, see Table1. The grey shading indicates antimicrobials classified as critically important (while the unshaded region indicates highly important antimicrobials).

Critically important antimicrobials in the quinolone drug class, like norfloxacin and ofloxacin, restricted in 2009 by the EU, experienced a higher prevalence of AMR in *Campylobacter* on organic farms in Europe compared to conventional farms: 91% (CI: 82-100%, n = 43) and 91% (CI: 82-100%, n = 43), respectively (Fig. 5C). In addition, resistance to ampicillin in *Campylobacter* in Europe was higher on organic farms (79%, CI: 67-91%, n = 43) than on conventional farms (66%, CI: 56-76%, n = 41; Fig. 5C). Similarly, in *Enterococcus* isolates, we found higher resistance to gentamicin on conventional farms in Oceania (66%, CI: 63-68%, n = 318). We also found lower resistance to ciprofloxacin in *Enterococcus* isolates on conventional farms in Europe 14% CI: 12-16%, n = 284 (Fig. 6B) and in North America 16% CI: 15-21%, n = 168 (Fig. 6C) as compared to Oceania which reported higher resistance prevalence of (84%, CI: 82-86%, n = 353 in *Enterococcus* isolates (Fig. 6D). The prevalence of AMR in *staphylococcus aureus* on conventional farms varied across regions. For instance, the level of resistance to cefoxitin was as high as 100% on conventional farms in Europe (n =36; Fig. 7A). Resistance to lincomycin in North America was 82% (CI: 73-87%, n=108) on conventional farms, while lincomycin resistance in Europe was much lower at 24% (CI: 17-33%, n =100; Fig. 7A and 7B).

**Figure 6.**
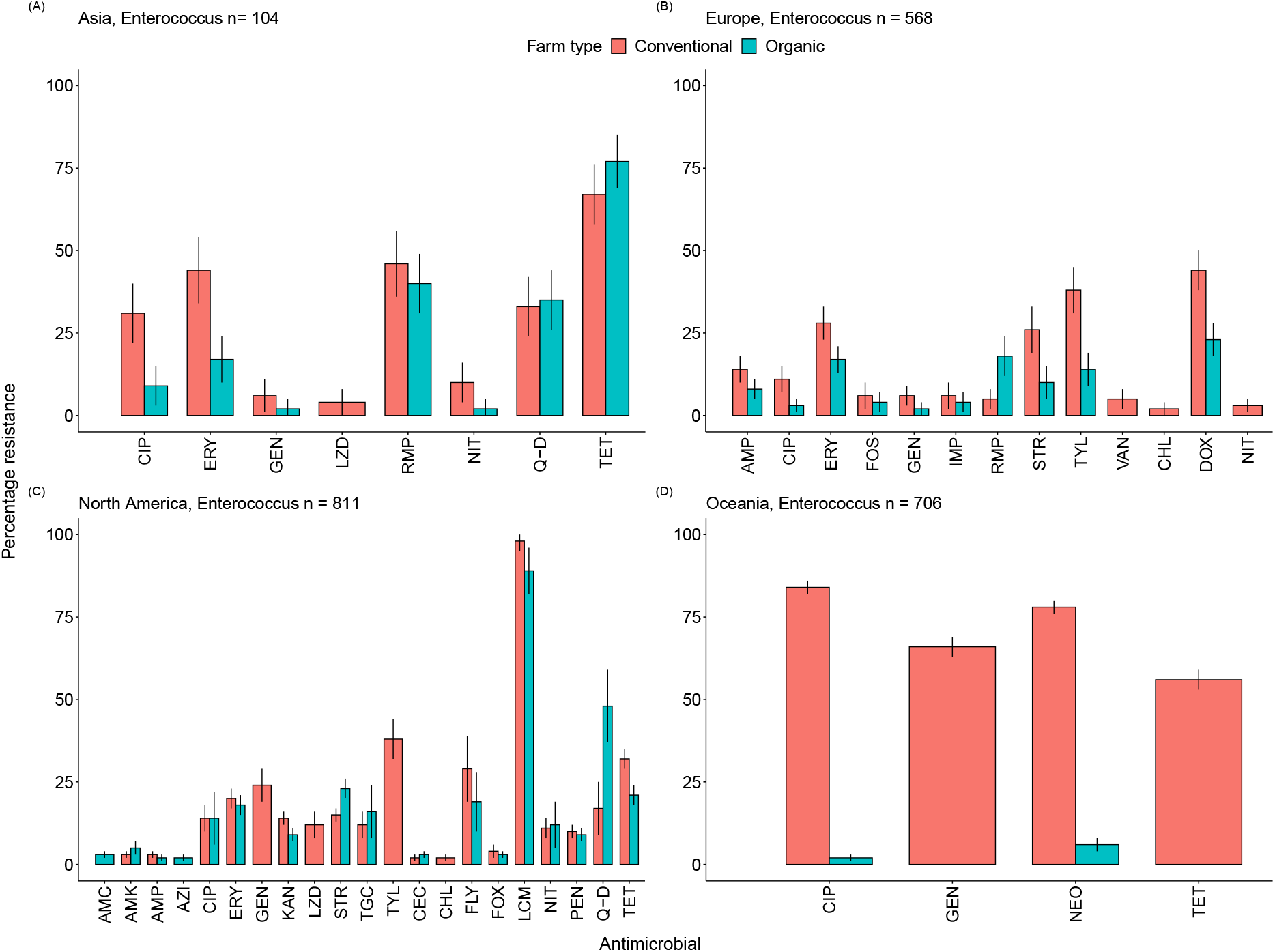
Patterns of antimicrobial resistance in *Enterococcus.* The percentage of antimicrobial resistance is shown for the number of isolates (n) examined on organic and conventional farms in each geographic region. A) Asia, *enterococcus,* n = 104, B) Europe, *enterococcus*, n = 568, C) North America, *enterococcus,* n = 811 and D) Oceania, *enterococcus,* n = 706. The error bars represent the 95% proportion confidence interval. For drug acronyms, see Table1. The grey shading indicates antimicrobials classified as critically important (while the unshaded region indicates highly important antimicrobials).

**Figure 7.**
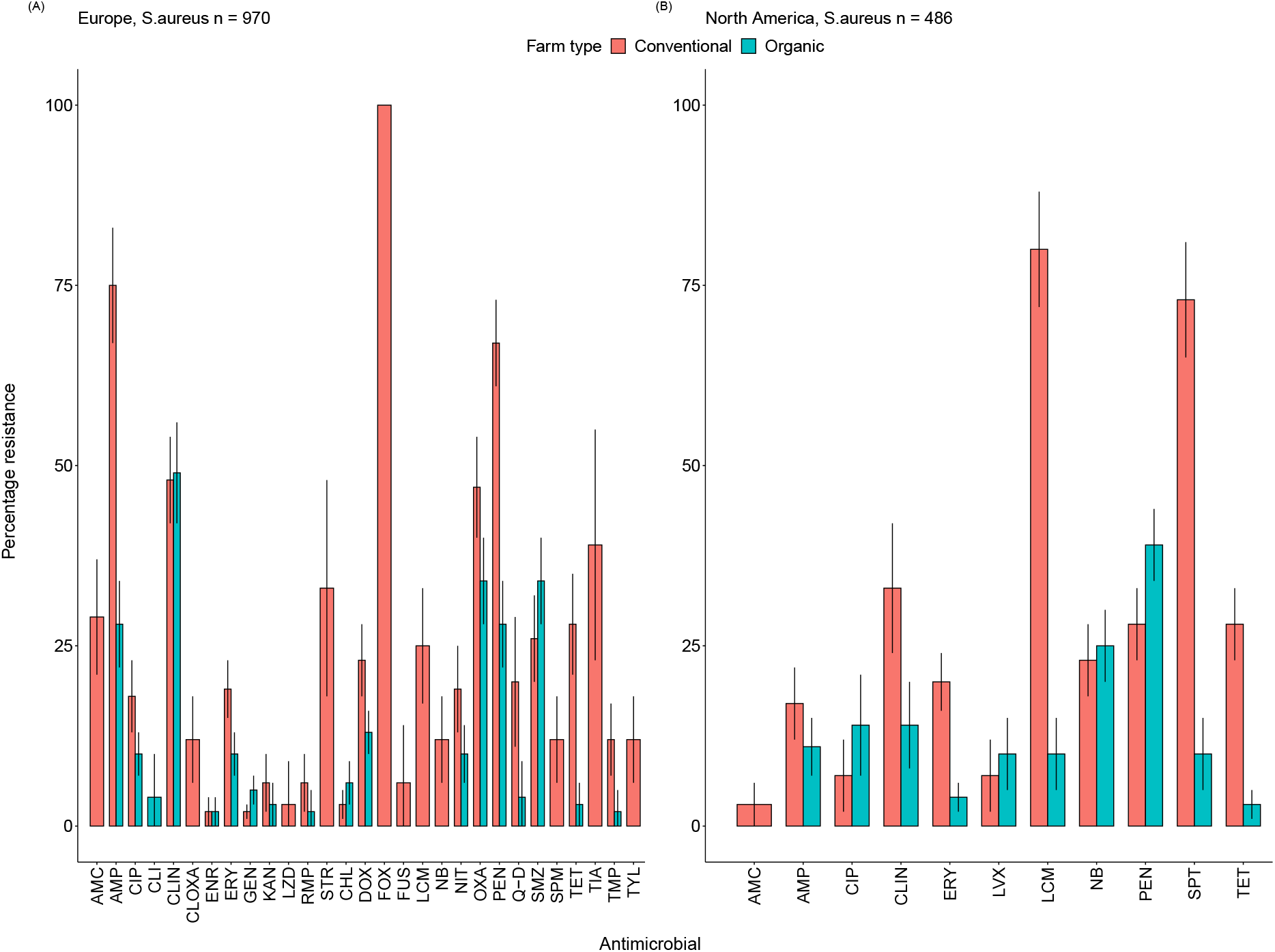
Patterns of antimicrobial resistance in *S. aureus.* The percentage of antimicrobial resistance is shown for the number of isolates (n) examined on organic and conventional farms in each geographic region. A) Europe, *S.aureus*, n = 970 and B) North America, *S.aureus*, n = 486. The error bars represent the 95% proportion confidence interval. For drug acronyms, see Table1. The grey shading indicates antimicrobials classified as critically important (while the unshaded region indicates highly important antimicrobials).

## DISCUSSION

Our findings reveal that overall, antimicrobial resistance (AMR) is lower on organic farms relative to their conventional counterparts. Specifically, organic farms had an AMR prevalence of 18%, compared to 28% on conventional farms. Moreover, our results suggest that between 2001 to 2020 AMR prevalence increased from 10-32% on organic farms and from 18-42% on conventional farms. The overall increase in AMR is alarming and suggests that the policies restricting drug use have had little overall success, due in part to loopholes in regulations and the global increase in consumer demand for meat products ^29^. For example, the Obama administration implemented policies to reduce antimicrobials as growth promoters in the US ^30^. This policy change, in turn, resulted in a shift in the labelling and marketing of these products from ‘growth promoters’ to ‘prophylactic therapeutics’ ^31^. In addition, global meat consumption doubled over the time frame covered here. This trend is expected to increase by 14% in 2030 ^32, 33^. This pattern is driven largely by improved economic security and growing urban populations in low and middle-income economies ^32^. Together, these contravening forces help clarify why the demand to reduce the use of antimicrobials in veterinary sectors has not resulted in a concomitant decrease in the prevalence of antimicrobial resistance in livestock.

Despite the overall lower prevalence of AMR on organic farms, we did find surprisingly high levels of AMR from *Campylobacter* isolates on organic farms. For instance, high resistance to ampicillin, a critically important antimicrobial, was reported in Europe (Fig. 5C). We also found higher AMR in *Campylobacter* from chicken isolates on organic farms in Portugal, the UK, and some parts of the US. These findings mirror previous research that has also found higher AMR in *Campylobacter* on chicken farms in the UK and Portugal ^34, 35^ and suggests that organic farming practices might not necessarily reduce consumer risk of exposure to *Campylobacter*.

*Campylobacter* poses a great threat to public health. For example, 60-80% of global *Campylobacter* cases originate from chicken products, and 400-500 million cases are reported globally every year ^36^, with 310,000 cases of potentially untreatable infections leading to 28 deaths annually in the US alone ‒ attributed to resistance to azithromycin and ciprofloxacin, two important anti-*Campylobacter* antibiotics ^19^. Therefore, eliminating this pathogen at the farm level will reduce the risk of human *Campylobacter* exposure and thus improve public health.

Surprisingly, we found higher AMR prevalence in environmental isolates from organic farms compared to conventional farms (Fig. 2C and 2D), which might have been a result of environmental contamination during the transition to organic farming. Specifically, these patterns were observed in parts of the US where most organic farms transition to conventional farms (Fig. 3). These findings highlight the role of the environment as a reservoir of antimicrobial resistance genes. In addition, we reported a high prevalence of resistance to norfloxacin and ofloxacin, drugs from the quinolone class in Europe (Fig. 5C). Previous studies have reported norfloxacin and ofloxacin among the top 90% of quinolones consumed by the EU countries ^37^. However, quinolone consumption has been restricted in the EU since then ^38^, thus calling for further investigations to better understand the impact of quinolone restrictions in Europe.

Our study highlights multidrug resistance (MDR) to drugs that are considered critically and highly important to human medicine in Asia, Europe, and North America on both organic and conventional farms (Fig. 4B, 5A, 5C, and 5D). Specifically, Europe reported multidrug resistance to drugs of the cephalosporin class in *E.coli*, while Asia reported multidrug resistance to drugs of the fluoroquinolone class in *Salmonella* on both organic and conventional farms.

These findings are consistent with previous research ^39–42^, which found high multidrug resistance in *E.coli* and *Salmonella*. Multidrug resistance has been linked to the misuse of antibiotics in animals and has led to failure of both cephalosporins and fluoroquinolones in treatment in Europe and Asia ^43–45^.

How do experimental patterns relate to broader usage patterns across countries? On the one hand, country-wide usage patterns are an admittedly coarse metric that lacks the requisite details to pinpoint the sectors of most concern. In the US, antimicrobial usage is not allowed on organic farms, while in Europe, according to the European regulation for organic dairy herds, a maximum of three treatments with antimicrobials are allowed per cow per year ^46–48^. On the other hand, mounting evidence indicates a strong correlation between drug use and the prevalence of antimicrobial resistance. Our results mirror these patterns. For instance, the highest prevalence of AMR was found in environmental isolates collected from conventional farms in the US (Fig. 3), a country that consistently ranks among the top three with high antimicrobial usage ^2, 49^. Conversely, countries like Sweden and New Zealand ^47, 50–52^ with low antimicrobial usage in food animals reported low AMR prevalence in the environment. Thus, while country-wide usage patterns are imperfect, they do provide an invaluable and robust metric for predicting and mitigating antimicrobial resistance.

Like other reviews, our study had limitations. To begin with, some continents were underrepresented. For instance, our review covered only one survey from South America and two from Oceania. Therefore, patterns from these continents should be interpreted with caution as they are likely to provide an inaccurate snapshot of actual resistance patterns. Additionally, our review included only studies written in English, which is an unfortunate but unavoidable omission that we hope to remedy in future studies. In addition, AMR point prevalence surveys pose a challenge to the standardization of data from susceptibility testing. Methodologies and protocols used in susceptibility testing may vary and result in inflation of AMR prevalence results. For example, in our study, the prevalence of AMR to spectinomycin and lincomycin was determined as a binary outcome (excluding intermediate resistance) ^28, 53^. This might have led to inflation of the results; therefore, results related to prevalence of resistance to spectinomycin and lincomycin presented in this study should be interpreted with caution.

The patterns presented here encompass a coarse snapshot that could be refined by more finely controlled experiments that account for at least three key variables. First, the studies included here likely differ in the timing of transition from conventional to organic farming. However, data on transition timing was not included in most studies, and therefore is beyond the scope of this analysis. Second, lack of global regulation standards for organic farming practices accounts for drastic differences in the time frame for implementation of restrictions on antimicrobial usage for animals on organic farms. For instance, the US National Organic Program standards allow for a one-year transition period from organic to conventional management in livestock systems and a three-year transition period from organic to conventional farming in crop systems ^54, 55^. Finally, the use of animal manure as fertilizer can increase drug residues in the environment and can select for higher antimicrobial resistance.

Additionally, animal manure can also contaminate the environment with antimicrobial resistance genes or plasmids. Recent advances in molecular and gene sequencing technologies have increased our awareness that these genes can transfer among bacteria as well as to other species, largely through horizontal gene transfer among microbes in the gut microbiome ^56–59^, leading to rapid transfer of multi-drug resistance in different hosts. The contaminated soil can function as a potential reservoir leading to an unexpectedly high AMR prevalence on organic farms ^59, 60^.

These factors underscore the need for an integrative approach, including restructuring policies, to help mitigate this public health problem and for further risk management investigations to clarify whether these patterns reflect environmental contamination or are related to a specific farming practice or pathogen outbreak.

To address these public health problems, techniques have been put in place to help reduce the risk of environmental contamination ‒ for instance, composting of organic manure, which happens through a biological process of anaerobic digestion involving humification and mineralization of organic matter. Compost resulting from the process by which environmental microorganisms break down organic materials helps eliminate or reduce pathogenic microorganism populations. The consequent lowering of AMR prevalence leads to economical, environmental, and public health advantages ^61, 62^.

We found evidence suggesting an increasing AMR trend on both farm types, as well as an alarming prevalence of AMR to some critically important antimicrobials. These trends are predicted to continue rising along with consumer demand for meat, which is projected to increase by 14% in 2030. Traditional interventions like stringent cleaning, antibiotics, and vaccines are critical for managing herd health and treating disease. In isolation, however, these costly and reactive approaches aimed at limiting pathogen proliferation concomitantly select for more virulent and resistant variants and ultimately ease their spread. The growing threat of antimicrobial resistance and consumer demands to reduce the use of antimicrobials in livestock emphasizes the need to leverage non-pharmacological approaches to prevent and manage disease. Organic farming practices hold promise in achieving this goal, but there is still room for improvement. Our results underscore the need for more concerted global efforts to develop unified organic, sustainable, and regenerative farming practices.

## Acknowledgements

We thank Dr. Eric Févre and Elizabeth Banset for helpful discussions that improved this manuscript.

## Author contribution

J.L.H and E.O.A designed the study. E.O.A and T.C performed the analysis. E.O.A and J.L.H collated the data. E.O.A, J.L.H, S.C.R and E.S wrote the manuscript. All authors contributed to revisions.

## Data availability statement

All data generated or analyzed during this study are included in this article and supplementary materials.

## Tables

**Table 1.** List of medically important antimicrobials, acronyms, antimicrobials classes and WHO classification included in the study.

| Antimicrobial | Antimicrobial compound | Antimicrobial class | WHO classification |
| --- | --- | --- | --- |
| Amikacin | AMK | Aminoglycoside | Critically important |
| Amoxicillin-clavulanic acid | AMC | Aminopenicillin - betalactamase inhibitors | Critically important |
| Ampicillin | AMP | Penicillin | Critically important |
| Ampicillin | AMP | Aminopenicillin - betalactamase inhibitors | Critically important |
| Apramycin | APR | Aminoglycoside | Critically important |
| Azithromycin | AZI | Macrolide | Critically important |
| Cefepime | CEP | 4th_gen cephalosporin | Critically important |
| Cefoperazone | CFP | 3rd_gen cephalosporin | Critically important |
| Cefotaxime | CTX | 3rd_gen cephalosporin | Critically important |
| Cefpodoxime | CPDX | 3rd_gen cephalosporin | Critically important |
| Ceftazadime | CAZI | 3rd_gen cephalosporin | Critically important |
| Ceftiofur | CTF | 3rd_gen cephalosporin | Critically important |
| Ceftiofur | CET | 3rd_gen cephalosporin | Critically important |
| Ceftiofur | CEF | 3rd_gen cephalosporin | Critically important |
| Ceftriaxone | CRO | 3rd_gen cephalosporin | Critically important |
| Ceftriaxone | CTX | 3rd_gen cephalosporin | Critically important |
| Cephalothin | CEF | 3rd_gen cephalosporin | Critically important |
| Ciprofloxacin | CIP | Quinolones | Critically important |
| Ciprofloxacin | CIP | Aminoglycoside | Critically important |
| Clindamycin | CLIN | Macrolide | Critically important |
| Cloxacillin | CLOXA | Penicillin | Critically important |
| Colistin | CST | Polymyxins | Critically important |
| Colistin | CTL | Polymyxins | Critically important |
| Enrofloxacin | ENR | Quinolones | Critically important |
| Erythromycin | ERY | Macrolide | Critically important |
| Fosfomycin | FOS | Phosphonic acid derivatives | Critically important |
| Gatifloxacin | GFX | Quinolones | Critically important |
| Gentamycin | GEN | Aminoglycoside | Critically important |
| Imipenem | IMP | Carbapenem | Critically important |
| Kanamycin | KAN | Aminoglycoside | Critically important |
| Levofloxacin | LVX | Quinolones | Critically important |
| Linezolid | LZD | Oxazolidinones | Critically important |
| Meropenem | MER | Carbapenem | Critically important |
| Mezlocillin | MZL | Penicillin | Critically important |
| Nalidixic acid | NAL | Quinolones | Critically important |
| Neomycin | NEO | Aminoglycoside | Critically important |
| Norfloxacin | NOR | Quinolones | Critically important |
| Ofloxacin | OFX | Quinolones | Critically important |
| Oxacillin | OXA | Penicillin | Critically important |
| Piperacilin | PIP | Penicillin | Critically important |
| Piperacilin | PIT | Penicillin | Critically important |
| Piperacillin-tazobactam | TZP | Penicillin | Critically important |
| Rifampicin | RMP | Ansamycin | Critically important |
| Spiramycin | SPM | Macrolide | Critically important |
| Streptomycin | STR | Aminoglycoside | Critically important |
| Tigecycline | TGC | Glycylcyclines | Critically important |
| Tylosin | TYL | Macrolide | Critically important |
| Vancomycin | VAN | Glycopeptides | Critically important |
| Amoxicillin-clavulanic acid | AMC | Aminopenicillin - betalactamase inhibitors | Highly important |
| Cefaclor | CEC | 2nd_gen cephalosporin | Highly important |
| Cefazolin | CFZ | 1st_gen cephalosporin | Highly important |
| Cefoxitin | FOX | 2nd_gen cephalosporin | Highly important |
| Cefoxitin | COX | 2nd_gen cephalosporin | Highly important |
| Ceftriaxone | CTX | 1st_gen cephalosporin | Highly important |
| Cefuroxime | CXM | 2nd_gen cephalosporin | Highly important |
| Cephalothin | CEF | 1st_gen cephalosporin | Highly important |
| Chloromphenicol | CHL | Amphenicol | Highly important |
| Chlortetracycline | CTET | Tetracycline | Highly important |
| Clindamycin | CLI | Lincosamides | Highly important |
| Doxycycline | DOX | Tetracycline | Highly important |
| Florfenicol | FLO | Amphenicol | Highly important |
| Florfenicol | FLL | Amphenicol | Highly important |
| Fusidic acid | FUS | Steroids antibacterials | Highly important |
| Lincomycin | LCM | Lincosamides | Highly important |
| Oxacillin | OXA | Penicillin | Highly important |
| Penicillin | PEN | Penicillin | Highly important |
| Quinopristin-dalfopristin | Q-D | Streptogramins | Highly important |
| Sulfamethaxazole | SMZ | Sulfonamide | Highly important |
| Sulfisoxazole | SMZ | Sulfonamide | Highly important |
| Tetracycline | TET | Tetracycline | Highly important |
| Trimethoprim | TMP | Sulfonamide | Highly important |
| Trimethoprim-Sulfamethaxazole | TMP-SMZ | Sulfonamide | Highly important |
| Nitrofurantoin | NIT | Nitrofurans derivatives | Important |
| Spectinomycin | SPT | Aminocyclitols | Important |
| Tiamulin | TIA | Pleuromutilins | Important |
| Flavomycin | FLY | Phosphoglycolipids | Not used in humans |
| Novobiocin | NB | Aminocoumarin |  |
| Penicillin-novobiocin | PEN-NB |  |  |

## Figure legends

## REFERENCES

1. Van Boeckel, T. P. et al. Global trends in antimicrobial resistance in animals in low- and middle-income countries. Science 365, eaaw1944 (2019).

2. Van Boeckel, T. P. et al. Global trends in antimicrobial use in food animals. Proc Natl Acad Sci U S A 112, 5649–5654 (2015).

3. Alimentarius, C. ORGANICALLY PRODUCED FOODS - CODEX ALIMENTARIUS. (2017).

4. Hosain, M. Z., Kabir, S. M. L. & Kamal, M. M. Antimicrobial uses for livestock production in developing countries. Vet World 14, 210–221 (2021).

5. Åkerfeldt, M. P., Gunnarsson, S., Bernes, G. & Blanco-Penedo, I. Health and welfare in organic livestock production systems—a systematic mapping of current knowledge. Org. Agr. 11, 105–132 (2021).

6. Rodrigues da Costa, M. & Diana, A. A Systematic Review on the Link between Animal Welfare and Antimicrobial Use in Captive Animals. Animals 12, 1025 (2022).

7. Willer, H., Trávníček, J., Meier, C. & Schlatter, B. <<The>> World of Organic Agriculture Statistics and Emerging Trends 2022. (Forschungsinstitut für biologischen Landbau FIBL, 2022).

8. Organic Trade Association. 2022 Organic Industry Survey Shows Steady Growth, Stabilizing Purchasing Patterns | OTA. https://ota.com/news/press-releases/22284 (2022).

9. Sapkota, A. R. et al. Lower prevalence of antibiotic-resistant Enterococci on U.S. conventional poultry farms that transitioned to organic practices. Environ Health Perspect 119, 1622–1628 (2011).

10. Mie, A. et al. Human health implications of organic food and organic agriculture: a comprehensive review. Environ Health 16, 111 (2017).

11. Gücükoğlu, A., Çadirci, Ö., Terzi Gülel, G., Uyanik, T. & Kanat, S. Organik Tavuk Etlerinden İzole Edilen Listeria monocytogenes İzolatlarinin Serotip ve Antibiyotik Direnç Profilinin Belirlenmesi. Kafkas Univ Vet Fak Derg (2020) doi:10.9775/kvfd.2019.23638.

12. Nunan, C. Report I Ending routine farm antibiotic use in Europe through improving animal health and welfare - EPHA. https://epha.org/ending-routine-farm-antibiotic-use/ (2022).

13. AccessScience Editors. U.S. bans antibiotics use for enhancing growth in livestock. (2017) doi:10.1036/1097-8542.BR0125171.

14. Alali, W. Q., Thakur, S., Berghaus, R. D., Martin, M. P. & Gebreyes, W. A. Prevalence and distribution of Salmonella in organic and conventional broiler poultry farms. Foodborne Pathog Dis 7, 1363–1371 (2010).

15. Quintana-Hayashi, M. P. & Thakur, S. Longitudinal study of the persistence of antimicrobial-resistant campylobacter strains in distinct Swine production systems on farms, at slaughter, and in the environment. Appl Environ Microbiol 78, 2698–2705 (2012).

16. Buntenkoetter, V. et al. Comparison of the phenotypic antimicrobial resistances and spa-types of methicillin-resistant Staphylococcus aureus (MRSA) isolates derived from pigs in conventional and in organic husbandry systems. Berl Munch Tierarztl Wochenschr 127, 135–143 (2014).

17. Hansson, I. et al. Differences in Genotype and Antimicrobial Resistance between Campylobacter spp. Isolated from Organic and Conventionally Produced Chickens in Sweden. Pathogens 10, 1630 (2021).

18. Murphy, C. P. et al. Scoping review to identify potential non-antimicrobial interventions to mitigate antimicrobial resistance in commensal enteric bacteria in North American cattle production systems. Epidemiol Infect 144, 1–18 (2016).

19. Yang, Y. et al. A Historical Review on Antibiotic Resistance of Foodborne Campylobacter. Front. Microbiol. 10, 1509 (2019).

20. Marchello, C. S., Carr, S. D. & Crump, J. A. A Systematic Review on Antimicrobial Resistance among Salmonella Typhi Worldwide. The American Journal of Tropical Medicine and Hygiene 103, 2518–2527 (2020).

21. Kijlstra, A. & Eijck, I. A. J. M. Animal health in organic livestock production systems: a review. NJAS: Wageningen Journal of Life Sciences 54, 77–94 (2006).

22. The EndNote Team. EndNote. (2013).

23. R core Team. R: The R Project for Statistical Computing. (2022).

24. QGIS Development Team. QGIS Geographic Information System. (2009).

25. Pinheiro, J. C., Bates, D. J., DebRoy, S. & Sakar, D. The Nlme Package: Linear and Nonlinear Mixed Effects Models, R Version 3. R package version vol. 6 (2012).

26. Burnham, K. P. & Anderson, D. R. Multimodel Inference: Understanding AIC and BIC in Model Selection. Sociological Methods & Research 33, 261–304 (2004).

27. Schweinberger, M. Fixed- and Mixed-Effects Regression Models in R. https://slcladal.github.io/regression.html (2022).

28. Schar, D. et al. Twenty-year trends in antimicrobial resistance from aquaculture and fisheries in Asia. Nat Commun 12, 5384 (2021).

29. Ritchie, H. & Roser, M. Meat and Dairy Production. Our World in Data (2017).

30. Jooma, S. Executive action to combat the rise of drug-resistant bacteria: is agricultural antibiotic use sufficiently addressed? J Law Biosci 2, 129–138 (2015).

31. Armbruster, W. J. & Roberts, T. The Political Economy of US Antibiotic Use in Animal Feed. in Food Safety Economics (ed. Roberts, T.) 293–322 (Springer International Publishing, 2018). doi:10.1007/978-3-319-92138-9_15.

32. OECD & Nations, F. and A. O. of the U. OECD-FAO Agricultural Outlook 2022-2031. (2022).

33. Statista. Global meat consumption by type 1990-2021. Statista

34. https://www.statista.com/statistics/274522/global-per-capita-consumption-of-meat/ (2022).

34. Carreira, A. C. et al. Comparative Genotypic and Antimicrobial Susceptibility Analysis of Zoonotic Campylobacter Species Isolated from Broilers in a Nationwide Survey, Portugal. Journal of Food Protection 75, 2100–2109 (2012).

35. Powell, L. F. et al. The prevalence of *Campylobacter* spp. in broiler flocks and on broiler carcases, and the risks associated with highly contaminated carcases. Epidemiol. Infect. 140, 2233–2246 (2012).

36. Igwaran, A. & Okoh, A. I. Human campylobacteriosis: A public health concern of global importance. Heliyon 5, e02814 (2019).

37. Adriaenssens, N. et al. Consumption of quinolones in the community, European Union/European Economic Area, 1997–2017. Journal of Antimicrobial Chemotherapy 76, ii37–ii44 (2021).

38. Kathrin Bauschab & Gernot Bonkatbc. Fluoroquinolone antibiotics – what we shouldn’t forget two years after the restriction by the European Commission. Swiss Med Wkly 151, (2021).

39. Melendez, S. N. et al. Salmonella enterica isolates from pasture-raised poultry exhibit antimicrobial resistance and class I integrons. J Appl Microbiol 109, 1957–1966 (2010).

40. Tanwar, J., Das, S., Fatima, Z. & Hameed, S. Multidrug Resistance: An Emerging Crisis. Interdisciplinary Perspectives on Infectious Diseases 2014, 1–7 (2014).

41. Muloi, D. et al. Epidemiology of antimicrobial-resistant Escherichia coli carriage in sympatric humans and livestock in a rapidly urbanizing city. International Journal of Antimicrobial Agents 54, 531–537 (2019).

42. Tawyabur, Md. et al. Isolation and Characterization of Multidrug-Resistant Escherichia coli and Salmonella spp. from Healthy and Diseased Turkeys. Antibiotics 9, 770 (2020).

43. Deguchi, T. et al. Fluoroquinolone Treatment Failure in Gonorrhea: Emergence of a: Neisseria gonorrhoeae: Strain With Enhanced Resistance to Fluoroquinolones. Sexually Transmitted Diseases 24, (1997).

44. Gagliotti, C., Buttazzi, R., Sforza, S. & Moro, M. L. Resistance to fluoroquinolones and treatment failure/short-term relapse of community-acquired urinary tract infections caused by Escherichia coli. Journal of Infection 57, 179–184 (2008).

45. Yang, J. et al. Occurrence and Characterization of Salmonella Isolated from Large-Scale Breeder Farms in Shandong Province, China. Biomed Res Int 2019, 8159567 (2019).

46. Sato, K., Bartlett, P. C. & Saeed, M. A. Antimicrobial susceptibility of Escherichia coli isolates from dairy farms using organic versus conventional production methods. J Am Vet Med Assoc 226, 589–594 (2005).

47. Sjöström, K. et al. Antimicrobial Resistance Patterns in Organic and Conventional Dairy Herds in Sweden. Antibiotics (Basel) 9, (2020).

48. US Department of Agriculture. Organic Regulations | Agricultural Marketing Service. https://www.ams.usda.gov/rules-regulations/organic (2022).

49. Tiseo, K., Huber, L., Gilbert, M., Robinson, T. P. & Van Boeckel, T. P. Global Trends in Antimicrobial Use in Food Animals from 2017 to 2030. Antibiotics (Basel) 9, E918 (2020).

50. Hillerton, J., Irvine, C., Bryan, M., Scott, D. & Merchant, S. Use of antimicrobials for animals in New Zealand, and in comparison with other countries. New Zealand Veterinary Journal 65, 71–77 (2017).

51. Khan, F. A., Söderquist, B. & Jass, J. Prevalence and Diversity of Antibiotic Resistance Genes in Swedish Aquatic Environments Impacted by Household and Hospital Wastewater. Front. Microbiol. 10, 688 (2019).

52. Pattis, I., Weaver, L., Burgess, S., Ussher, J. E. & Dyet, K. Antimicrobial Resistance in New Zealand-A One Health Perspective. Antibiotics (Basel) 11, 778 (2022).

53. Rhodes, S. et al. Getting ahead of antibiotic-resistant Staphylococcus aureus in U.S. hogs. Environmental Research 196, 110954 (2021).

54. Oregon tilth. Converting livestock to organic. Oregon Tilth https://tilth.org/knowledgebase_category/converting-livestock/ (2019).

55. USDA. Organic Transitioning | Agricultural Marketing Service. https://www.ams.usda.gov/services/organic-certification/transitioning-to-organic (2019).

56. Shterzer, N. & Mizrahi, I. The animal gut as a melting pot for horizontal gene transfer. Can. J. Microbiol. 61, 603–605 (2015).

57. Gurmessa, B. et al. Variations in bacterial community structure and antimicrobial resistance gene abundance in cattle manure and poultry litter. Environmental Research 197, 111011 (2021).

58. Algarni, S., Ricke, S. C., Foley, S. L. & Han, J. The Dynamics of the Antimicrobial Resistance Mobilome of Salmonella enterica and Related Enteric Bacteria. Front. Microbiol. 13, 859854 (2022).

59. Vinayamohan, P. G., Pellissery, A. J. & Venkitanarayanan, K. Role of horizontal gene transfer in the dissemination of antimicrobial resistance in food animal production. Current Opinion in Food Science 47, 100882 (2022).

60. Hiltunen, T., Virta, M. & Laine, A.-L. Antibiotic resistance in the wild: an eco-evolutionary perspective. Phil. Trans. R. Soc. B 372, 20160039 (2017).

61. Collignon, P. & McEwen, S. One Health—Its Importance in Helping to Better Control Antimicrobial Resistance. TropicalMed 4, 22 (2019).

62. Lima, T., Domingues, S. & Da Silva, G. J. Manure as a Potential Hotspot for Antibiotic Resistance Dissemination by Horizontal Gene Transfer Events. Vet Sci 7, E110 (2020).

