## Supplementary note 1-3 for "Global trends in antimicrobial resistance on organic and conventional farms"

This document contains:

1. Supplementary note1
2. Supplementary note 2
3. Supplementary note 3

Supplementary note1: Literature search

We searched three databases (PubMed, Web of Science and PubAg) for point prevalence surveys and extracted data on antimicrobial resistance in livestock organic and conventional farms in Oceania, South America, North America, Europe and Asia. All the surveys were published between 2000 and 2020 in the English language.

We used the following unique formulas for each database:

a) PubMed search terms:

**Livestock:**

( "goats" [MeSH Terms] OR "goat" [TW] OR "goats" [TW] OR "capra" [TW] OR "caprine" [TW] OR "caprines" [TW] OR "cattle" [MeSH Terms] OR "cow" [TW] OR "cows" [TW] OR "cattle" [TW] OR "bovine" [TW] OR "bos" [TW] OR "sheep, domestic" [MeSH terms] OR "sheep" [TW] OR "lamb" [TW] OR "ovis aries" [TW] OR "poultry" [MeSH Terms] OR "poultry" [TW] OR "turkey" [TW] OR "chicken" [TW] OR "duck" [TW] OR "geese" [TW] OR "domestic fowl" [TW] OR "Sus scrofa" [MeSH terms] OR "swine" [TW] OR "sow" [TW] OR "pig" [TW] OR "pigs" [TW] OR "livestock" [MeSH Terms] OR ("livestock" [TW] AND "meat" [TW]) OR ("livestock" [TW] AND ("dairy" [TW] or "milk" [TW])) OR "beef" [TW] OR "meat production" [TW] OR "dairy production" [TW] OR "dairy farm*" [TW] OR pork [TW] OR ("broiler" [TW] AND ("chick" [TW] OR "flock" [TW])) OR ("layer" [TW] AND ("chick" [TW] OR "flock" [TW])) OR "egg" [TW] OR "eggs" [TW] OR "cheese" [TW] OR "milk" [TW] OR "cheese" [MeSH Terms] OR "Milk" [MeSH Terms] OR "cultured milk products" [MeSH Terms] )

**Bacteria:**

( "campylobacter"[MeSH Terms] OR "campylobacter"[TW] OR "campylobacters"[TW] OR "escherichia coli"[MeSH Terms] OR "e coli"[TW] OR "escherichia coli"[TW] OR "salmonella"[MeSH Terms] OR "salmonella"[TW] OR "salmonellas"[TW] OR "salmonellae"[TW] OR "staphylococcus"[MeSH Terms] OR "staphylococcus"[TW] OR "staphylococcu"[TW] OR "methicillin resistant staphylococcus aureus" [TW] OR "MRSA"

[TW] OR "staphylococcus aureus" [TW] OR "enterococcus"[MeSH Terms] OR "enterococcus"[TW] OR "enterococcu"[TW])

**Antibiotic Resistance:**

( "drug resistance, microbial" [MeSH terms] OR ("antimicrobial" [TW] AND "resistan*" [TW]) OR ("antibacterial" [TW] AND "resistan*" [TW]) OR ("antibiotic" [TW] AND "resistan*" [TW]) OR (("drug" [TW] AND "resistan*" [TW]) AND (antibiotic [tw] OR antibiotics[tw])) OR "antimicrobial susceptibility" [TW] OR "antibacterial susceptibility" [TW] OR "antibiotic susceptibility testing" [TW] OR "antimicrobial susceptibility patterns" [TW] OR "antimicrobial stewardship" [MeSH Terms] OR "antimicrobial stewardship" [TW] OR "microbiological profile" [TW] OR "microbiological profiles" [TW] OR "microbiological profiling" [TW] OR "phylogenetic profile" [TW] OR "phylogenetic profiles" [TW] OR "phylogenetic profiling" [TW] )

**Organic/Conventional Farming:**

( "agriculture" [MeSH Terms] OR ("organic" [TW] AND "agriculture" [TW]) OR ("organic" [TW] AND "farm*" [TW]) OR ("conventional" [TW] AND "agriculture" [TW]) OR ("conventional" [TW] AND "farm*" [TW]) OR (("agriculture" [TW] OR "farm*") AND ("with antibiotic*" OR "without antibiotic*" OR "antimicrobial free")) OR "organic livestock" [TW] OR ("organic" [TW] AND "dair*" [TW]) OR ("conventional" [TW] AND "dair*" [TW]) OR ("organic" [TW] AND "flock*" [TW]) OR ("conventional" [TW] AND "flock*" [TW]) OR ("organic" [TW] AND "herd*" [TW]) OR ("conventional" [TW] AND "herd*" [TW]) )

b) Web of Science search terms:

**Livestock:**

livestock OR cattle OR cow* OR bovine* OR beef OR herd OR milk OR cheese OR sheep* Or lamb* OR goat* OR pig* OR swine OR Sow OR meat OR pork OR chick* OR flock OR poultry OR egg* OR broiler OR turkey* OR geese

**Bacteria:**
bacteri* OR "Escherichia coli" OR "E. Coli" OR Salmonella* OR Staphylococcus OR MRSA OR Campylobacter OR Enterococcus OR "enteric bacteria"

**Antibiotic Resistance:**
(antimicrob* NEAR/2 resistan*) OR (antibiotic* NEAR/2 resistan*) OR (drug NEAR/2 resistan*) OR (antibacter* NEAR/2 resistan*) OR (antimicrob* NEAR/2 susceptib*) OR (antibacter* NEAR/2 susceptib*) OR (antibiotic NEAR/2 susceptib*)

**Organic/conventional farming:**
(conventional NEAR/2 farm*) OR "conventional agriculture" OR (farm NEAR/2 antibiotic*) OR (farm NEAR/2 antimicrob*) OR (conventional NEAR/2 livestock) OR (conventional NEAR/2 dairy) OR (conventional NEAR/2 poultry) OR (conventional NEAR/2 egg*) OR (organic NEAR/2 farm*) OR (organic NEAR/2 agriculture) OR (farm NEAR/2 antibiotic*) OR (organic NEAR/2 livestock) OR (organic NEAR/2 dairy) OR (organic NEAR/2 poultry) OR (organic NEAR/2 egg*)

c) Pub Ag search terms:

((antimicrobial OR antibiotic OR drug) AND (resistance OR resistant OR susceptibility)) AND ((conventional OR organic) AND (agriculture OR farming OR dairy OR livestock OR poultry))

d) Grey literature search terms:

(Antimicrobial resistance OR antibiotic resistance OR drug resistance) AND (bacteria species) AND (conventional* OR organic*)

We did not add geographic location as a search parameter because geographic location often does not work well as a keyword.

The grey literature searched includes:

a) National Antibiotic Monitoring Program (USDA)
b) NARMS Reports/Summaries | NARMS interactive data
c) FAO Antimicrobial Resistance

d) WHO GLASS database
e) ECDC/EFSA/EMA first joint report on the integrated analysis of the consumption of antimicrobial agents and occurrence of antimicrobial resistance in bacteria from humans and food-producing animals
f) Antimicrobial resistance surveillance in Europe 2022 - 2020 data

Reviews, book chapters, unrelated topics, non-English records and meta-analyses were excluded. As a way of assessing data quality in our review, we excluded records where farm types were not clearly identified as organic or conventional, no geographic information on study location was given, resistance rates were unclear and studies involving imported products. Data extraction results were stratified according to the country name, antimicrobial resistance results, farm type, and pathogens

Supplementary note 2: Model Diagnostics

Assumptions

Linear mixed effects models are based on the following assumptions (Schweinberger, 2022):

1. Independent variables measured without an error (Fixed x)
2. Errors are normally distributed
3. Errors are independent
4. Errors have constant variance
5. The relationship between X and Y is linear

We used the graphs to determine whether the model assumptions are met

a) Turkey-Anscombe plot

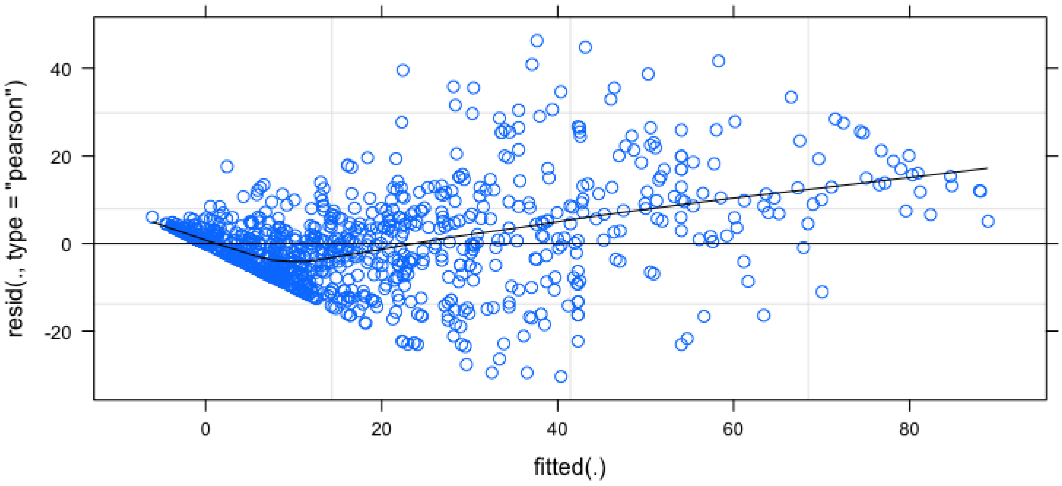


The above figure, demonstrates homoscedasticity (stable variance) meaning the variance of the residuals does not seem to increase or decrease with the fitted values

b) Quantile-quantile (Q-Q) plots

We assumed that both the errors and the random intercepts come from a normal distribution and used Q-Q plot to test this.

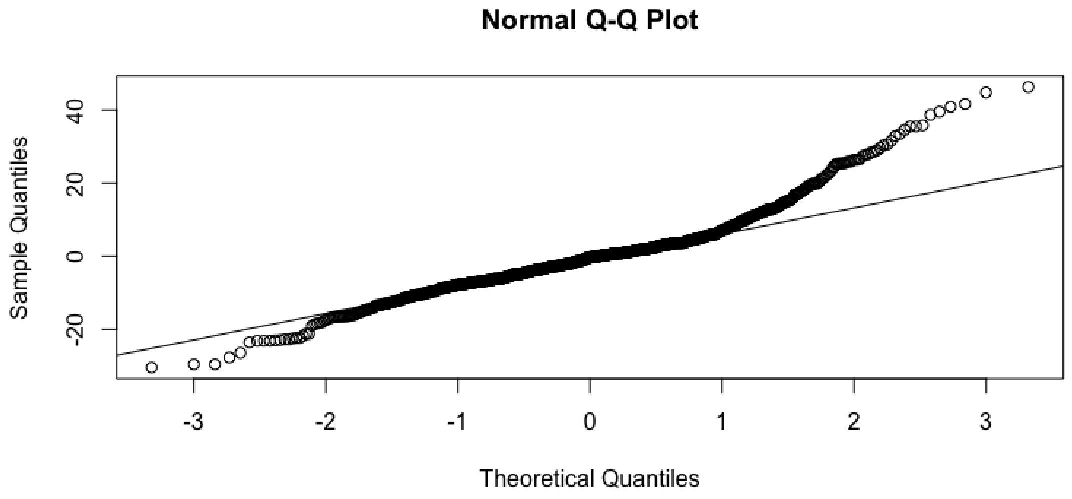


The QQ-plot above shows that there is no clear evidence that the our data does not follow a normal distribution. Although not perfect, this plot does not show a clear violation of the normality assumption for the random effects.

c) Residuals against predictors

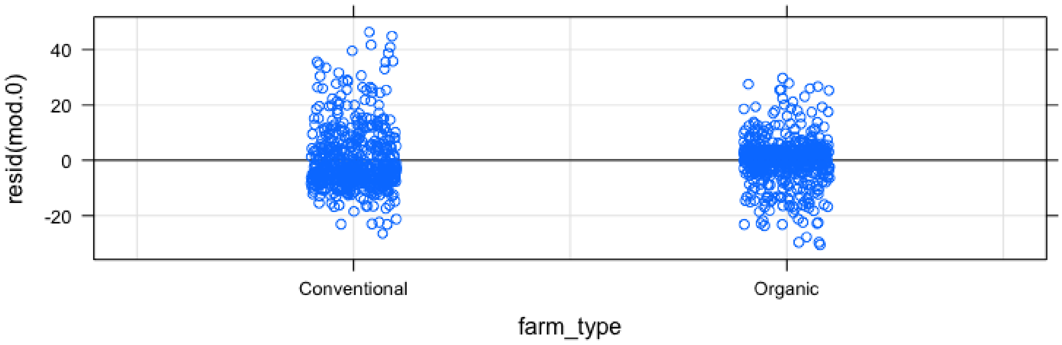


The figure above indicates that residuals on conventional farms have a higher variance as compared to organic farms, nevertheless, the is no dramatic difference between the two farm types.

To better identify deviations from linearity we add a line connecting the average values at each host. We then plot the residuals against the host.

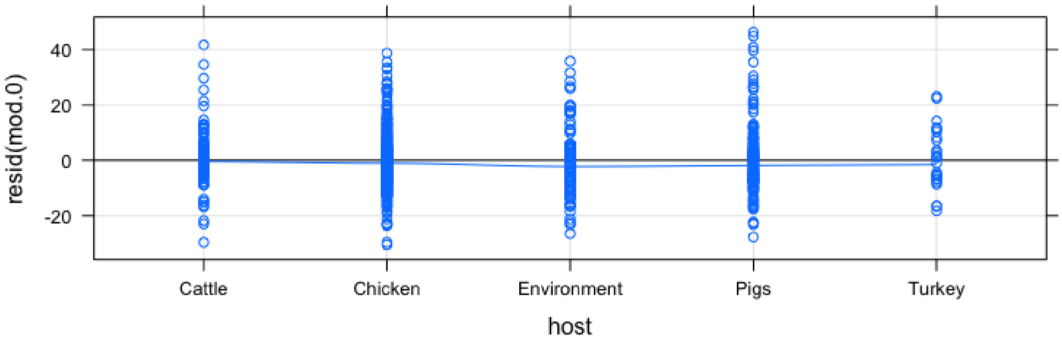


There seems to be no clear deviation from zero. This implies that at this stage there is no evidence against percent resistance of modelled hosts. Moreover, the variance of the residuals seems to be constant along the five-time points (hosts) (i.e homoscedasticity).

We used the dot plot below to check whether all studies have similar variances.

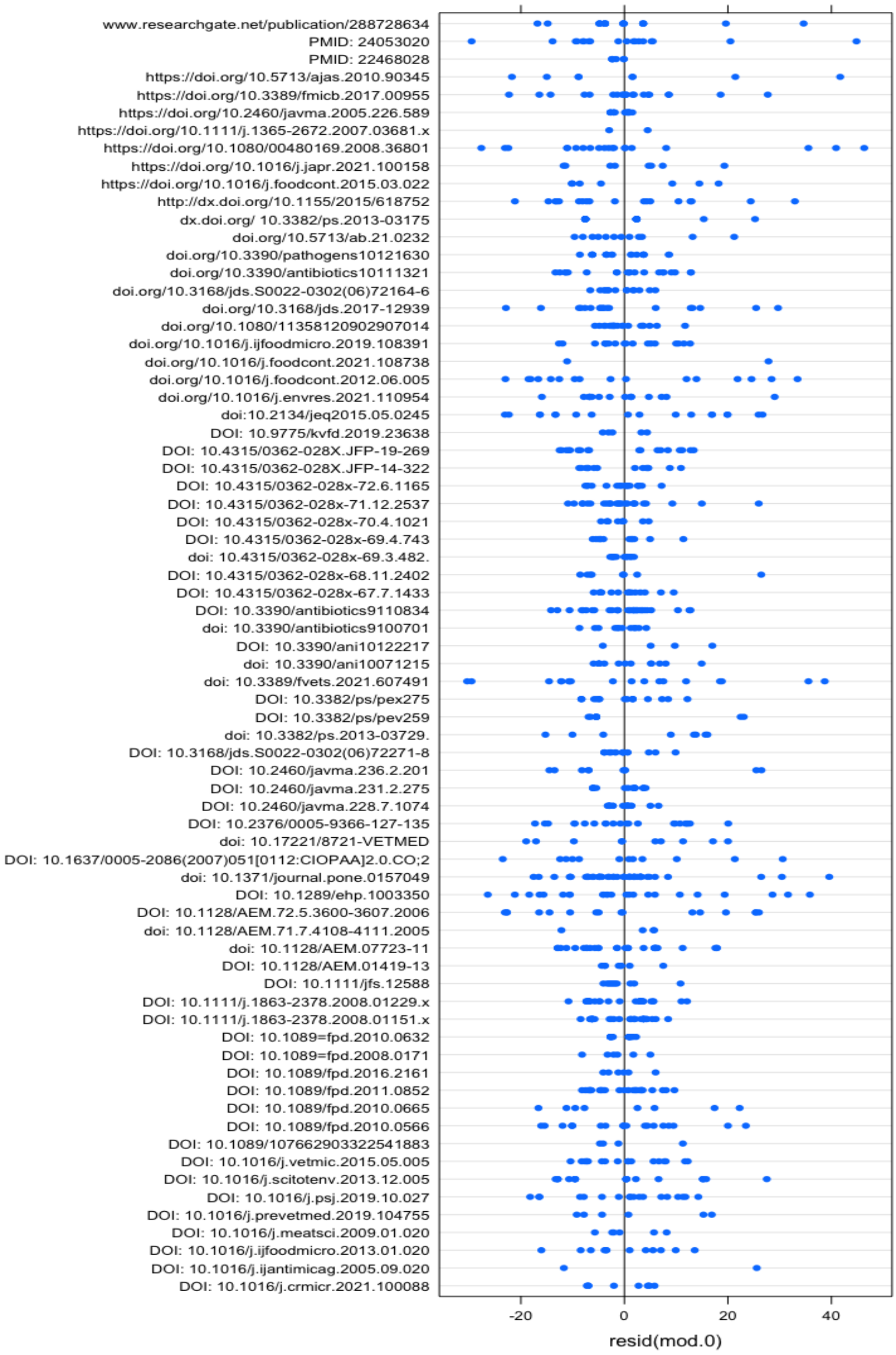


The above dot plot shows that there is no striking difference in variability among the studies.

**REFERENCES**

Schweinberger, M. (2022) *Fixed- and Mixed-Effects Regression Models in R*. Available at: https://slcladal.github.io/regression.html (Accessed: 11 October 2022).

### Supplementary note 3: Antimicrobial class prevalence in chicken by country

The graph below represents the antimicrobial resistance prevalence of different antimicrobial classes in chickens across different countries.

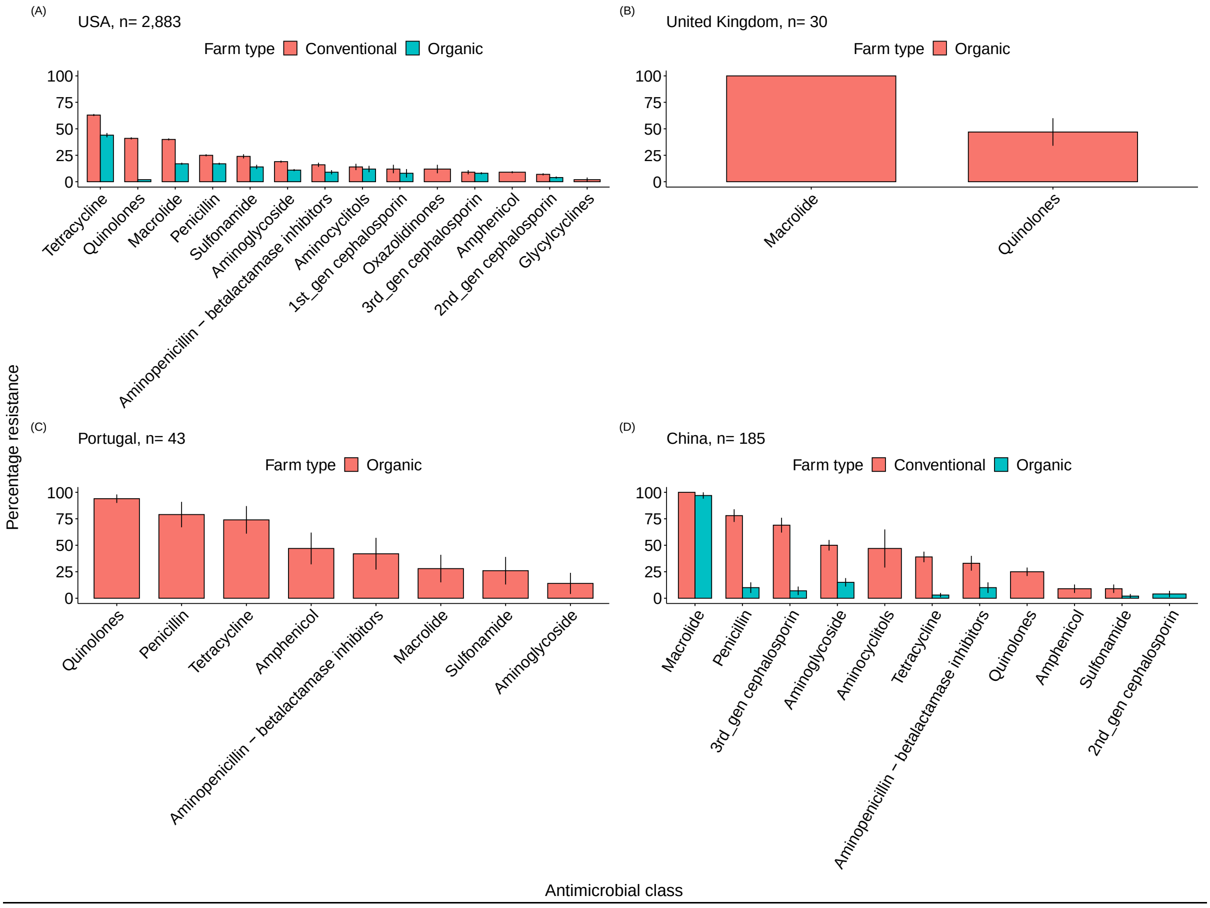


**Patterns of antimicrobial resistance in chicken.** The percentage of antimicrobial resistance is shown for the number of isolates (n) examined on organic and conventional farms in each country. Error bars represent the 95% proportion confidence interval.
